# Differential contributions of left-hemispheric language regions to basic semantic composition

**DOI:** 10.1101/2019.12.11.872457

**Authors:** Astrid Graessner, Emiliano Zaccarella, Gesa Hartwigsen

**Affiliations:** Lise-Meitner Research Group Cognition and Plasticity, Max Planck Institute for Human Cognitive and Brain Sciences, Leipzig, Germany; Department of Neuropsychology, Max Planck Institute for Human Cognitive and Brain Sciences, Leipzig, Germany

**Keywords:** Meaning composition, angular gyrus, conceptual combination, fMRI, functional connectivity

## Abstract

Semantic composition, i.e. the ability to combine single words to form complex meanings, is a core feature of human language. Despite growing interest in the basis of semantic composition, the neural correlates and the interaction of regions within this network remain a matter of debate. In the present fMRI study, we designed a well controlled two-word paradigm in which phrases only differed along the semantic dimension while keeping syntactic information alike. 33 healthy participants listened to meaningful phrases (“fresh apple”), anomalous phrases (“awake apple”) and pseudoword phrases (“awake gufel”) while performing both an implicit and an explicit semantic task. We identified neural signatures for distinct processes during basic semantic composition: The *general phrasal composition* process, which is independent of the plausibility of the resulting phrase, engages a wide-spread left hemispheric network comprising both executive semantic control regions as well as general conceptual representation regions. Within this network, the functional connectivity between the left anterior inferior frontal gyrus, the bilateral pre-supplementary motor area and the posterior angular gyrus specifically increases during *meaningful phrasal composition*. The anterior angular gyrus, on the other hand, guides more *specific compositional* processing evaluating phrasal plausibility. Overall, our results were stronger in the explicit task, pointing towards partly task-dependent involvement of the regions. Here we provide a separation between distinct nodes of the semantic network, whose functional contributions depend on the type of compositional process under analysis. For the first time, we show that the angular gyrus may be decomposable into two sub-regions during semantic composition.

## Introduction

One of the core features of human language is the ability to combine single words into complex phrases. Semantic knowledge helps us making sense of individual words and semantic composition processes drive the way we combine individual meanings into more composite ones. Current neuroanatomical models of semantic processing highlight a widely distributed fronto-temporo-parietal network in the left hemisphere (Binder et al., 2009). Focusing on semantic composition, a number of studies have identified several brain regions in the left hemisphere showing higher activation for sentences than word lists, including the angular gyrus (AG) in the inferior parietal lobe, posterior middle temporal gyrus (pMTG), anterior temporal lobe (ATL) and the anterior inferior frontal gyrus (aIFG) (Brennan & Pylkkänen, 2012; Humphries et al., 2006; Lau et al., 2008; Matchin et al., 2017; Pallier et al., 2011; Vandenberghe et al., 2002; Vigneau et al., 2006). However, due to the complexity of sentential manipulations, it can be difficult to differentiate processes focusing on semantic composition from those including other cognitive domains such as syntax, working memory, attention and cognitive control (Badre, 2008; Makuuchi & Friederici, 2013). In recent years, neuroscientific researchers have become increasingly more interested in simpler paradigms using two- or three-word phrases to tackle semantic composition in more controlled linguistic constructions (Pylkkänen, 2019).

One of the proposed key regions for basic semantic composition is the AG. Several studies report recruitment of the left (and sometimes right) AG for two-word phrases relative to single words (Bemis & Pylkkänen, 2013b), meaningful as compared to meaningless adjective-noun phrases (Graves et al., 2010; Molinaro et al., 2015; Price et al., 2015) and for tracking thematic relations between words (Boylan, Trueswell, & Thompson-Schill 2015; Boylan, Trueswell, & Thompson-Schill 2017; Lewis, Poeppel, & Murphy 2018).

Another region that has consistently been implicated as a key semantic composition region is the ATL. Within the “hub-and-spokes” model, the ATL is considered to bind information from different modalities (the “spokes”), serving as a transmodal “hub” (see Lambon Ralph et al. 2016). This view is supported by the finding that patients with semantic dementia who show bilateral atrophy of the ATL are impaired in semantic processing across all input modalities and types of concepts (Mummery et al., 2000). Regarding basic semantic composition, most evidence for the ATL as the key region of conceptual combination is derived from Magnetoencephalography (MEG) studies. Increased activity in the bilateral ATL has been shown for two-word phrases as compared to single words in both the visual and auditory modality (Bemis & Pylkkänen, 2013b) and in different languages, including English, Spanish, Arabic, and American Sign Language (Bemis & Pylkkänen, 2011; Blanco-Elorrieta et al., 2018; Molinaro et al., 2015; M. Westerlund et al., 2015). Other studies, however, also found that ATL activity is stronger when composition leads to an increase in the specificity of the concept (e.g. higher ATL activation for “tomato dish” than “vegetable dish”) (Zhang & Pylkkänen, 2015).

Functional MRI evidence for the ATL as key region for basic semantic composition is rather scarce, which might be explained by several methodological issues. First, the ATL suffers from signal loss in fMRI due to its location near the sinuses. Secondly, the use of low-level baselines which still engage semantic processing (e.g., internal speech) might have led to a lower likelihood of finding ATL activation because semantic activation in this region was removed during the subtraction analyses (Visser et al., 2010).

Finally, a region that has received less attention in the context of semantic composition but has rather been referred to as an executive semantic control region is the left aIFG (BA45/47). Increased aIFG activation has been observed in previous studies for sentences compared to word lists (Matchin, Liao, et al., 2019; Pallier et al., 2011), ambiguous relative to unambiguous sentences (Vitello et al., 2014) and two-word phrases compared to single words (Schell et al., 2017). The latter authors interpret the role of BA45 in the context of basic semantic composition as tracking the amount of words that can be integrated into context. Thus, two-word phrases should always elicit higher activity in left aIFG than single words.

Despite considerable effort to characterize the neural correlates for basic semantic composition, several questions remain open. First, although there is consensus that left AG, ATL and aIFG play important roles in semantic composition, most fMRI studies have focused on the contribution of single regions instead of investigating functional interactions at a larger network level. Consequently, it remains unclear how these regions influence each other during successful composition. At the single-word level, Hartwigsen et al. (2015) have shown that left AG and aIFG were able to compensate for a focal disruption of the respective other region induced by transcranial magnetic stimulation (TMS). In that study, semantic decisions were only impaired after both regions had been perturbed, showing that the interplay of these regions is causally relevant for semantic decisions. However, it is not clear whether this interaction is restricted to the single-word level or whether it is also involved in compositional processing.

Furthermore, it remains unclear whether the previously reported recruitment of brain areas during semantic composition is task-dependent. While some studies explicitly asked subjects to compose the meaning of the stimuli (Bemis & Pylkkänen, 2011; Graves et al., 2010; Price et al., 2015; Schell et al., 2017), others intentionally did not (Graves et al., 2010; Matchin, Brodbeck, et al., 2019; Matchin et al., 2017; Matchin, Liao, et al., 2019; Molinaro et al., 2015). Few studies have directly compared different tasks while keeping the stimulus material similar and different results between studies might thus reflect differences in task processing and demand.

Finally, we note that the few existing two-word studies have used different baseline conditions. While some studies compared the processing of two-word phrases to single words (e.g., Bemis & Pylkkänen, 2011, 2013a; Schell et al., 2017), others looked at more meaningful versus less meaningful two-word combinations (Graves et al., 2010; Molinaro et al., 2015; Price et al., 2015). It is conceivable that the different semantic regions discussed above fulfill distinct tasks in the semantic composition process such as tracking the amount of words versus combining the meaning of two separate concepts into a whole. To address this issue, it is thus advantageous to investigate the potential different processes in one single experimental setting.

In the current experiment, we created a paradigm consisting of three different two-word phrases: meaningful phrases (“fresh apple”), anomalous phrases (“awake apple”), and pseudoword phrases containing pseudonouns (“awake gufel”; see Methods section for details). This design allowed us to first look into the semantic processes guiding the composition of more complex meanings compared to simpler ones. Thus, we could separate (1) *meaningful phrasal composition* (meaningful > pseudoword phrase) from (2) *anomalous phrasal composition* (anomalous > pseudoword phrase). Additionally, the overlap of the two contrasts would show regions that are involved in (3) *general phrasal composition*, independent of the plausibility of the resulting phrase. Second, given that the syntactic information was kept constant across conditions, we could further measure two more *specific processes* directly tackling semantic plausibility: (4) *specific meaningful composition* (meaningful > anomalous) and (5) *specific anomalous composition* (anomalous > meaningful). Further, to explore whether the activation of semantic core regions is task-dependent, we included both an implicit and an explicit semantic task. This allowed us to distinguish task-specific and automatic processes during semantic composition. If semantic composition occurs implicitly and independent of task demands, we should see similar results for the explicit and implicit tasks.

Finally, to assess task-related changes in functional connectivity within the semantic composition network, we conducted psychophysiological interaction analyses. This network perspective has grown increasingly more popular in the study of neurocognitive processes as it has become clear that the brain is organized in large scale networks (Bassett & Sporns, 2017; Hartwigsen, 2018). In summary, we aimed to provide a comprehensive characterization of the network for basic semantic composition and explore potential task dependencies in the activation patterns.

Based on the above-cited studies, we expected to find high involvement of the left AG, ATL and aIFG during *meaningful phrasal composition*, reflecting semantic processes guiding the formation of complex meanings from simpler ones. Within this network, the AG and the ATL should be more specifically recruited during the processing of meaningful phrases compared to anomalous phrases (*specific meaningful composition*) as an effect of semantic plausibility, regardless of phrasal complexity. Conversely, the aIFG should be maximally recruited during the processing of anomalous phrases compared to meaningful phrases (*specific anomalous composition*), as a function of higher semantic control. By administering both an implicit and an explicit task, we aimed to identify regions that are activated in a task-dependent manner. We expected more inferior frontal involvement for the explicit task, while we hypothesized regions that are associated with automatic semantic processing (e.g. left ATL, AG) to be activated also in the implicit task. A potential overlap in activation for the two tasks was expected to reflect task-independent semantic composition processes. Regarding functional connectivity, we hypothesized that an interaction between left AG and aIFG, as previously observed during single word processing, would drive the comprehension of two-word phrases. However, we were also interested in the connectivity of other semantic network regions and therefore conducted several exploratory analyses with seed regions from the univariate GLM results.

## Methods

### Participants

Thirty-seven right-handed German-speaking subjects participated in the two sessions of the study. They had normal hearing, corrected to normal vision and no history of neurological disorders or contraindication to MR-scanning. Four participants had to be removed from the analyses due to low task accuracy (see Behavioral Analysis). The final group of participants that entered the analyses consisted of 33 participants (16 females, mean age 26 years, SD = 3.6 years).

All participants gave their written informed consent and were reimbursed with 10€/hour. The study protocol conformed to the principles of the Declaration of Helsinki and was approved by the local ethics committee at the University of Leipzig.

### Experimental Paradigm

All participants completed two event-related fMRI sessions separated by at least one week with two different tasks on the same set of stimuli. Auditory stimuli were presented using the software package Presentation (Neurobehavioral Systems, Inc., Albany, CA, USA) via MR-compatible in-ear headphones (MR-Confon, Magdeburg, Germany). Volume was adjusted to an optimal individual hearing level. Stimuli consisted of spoken word pairs that were either meaningful (“fresh apple”), anomalous (“awake apple”) or had the noun replaced by a pseudoword (“awake gufel”) (see Figure 1A).

**Figure 1.**
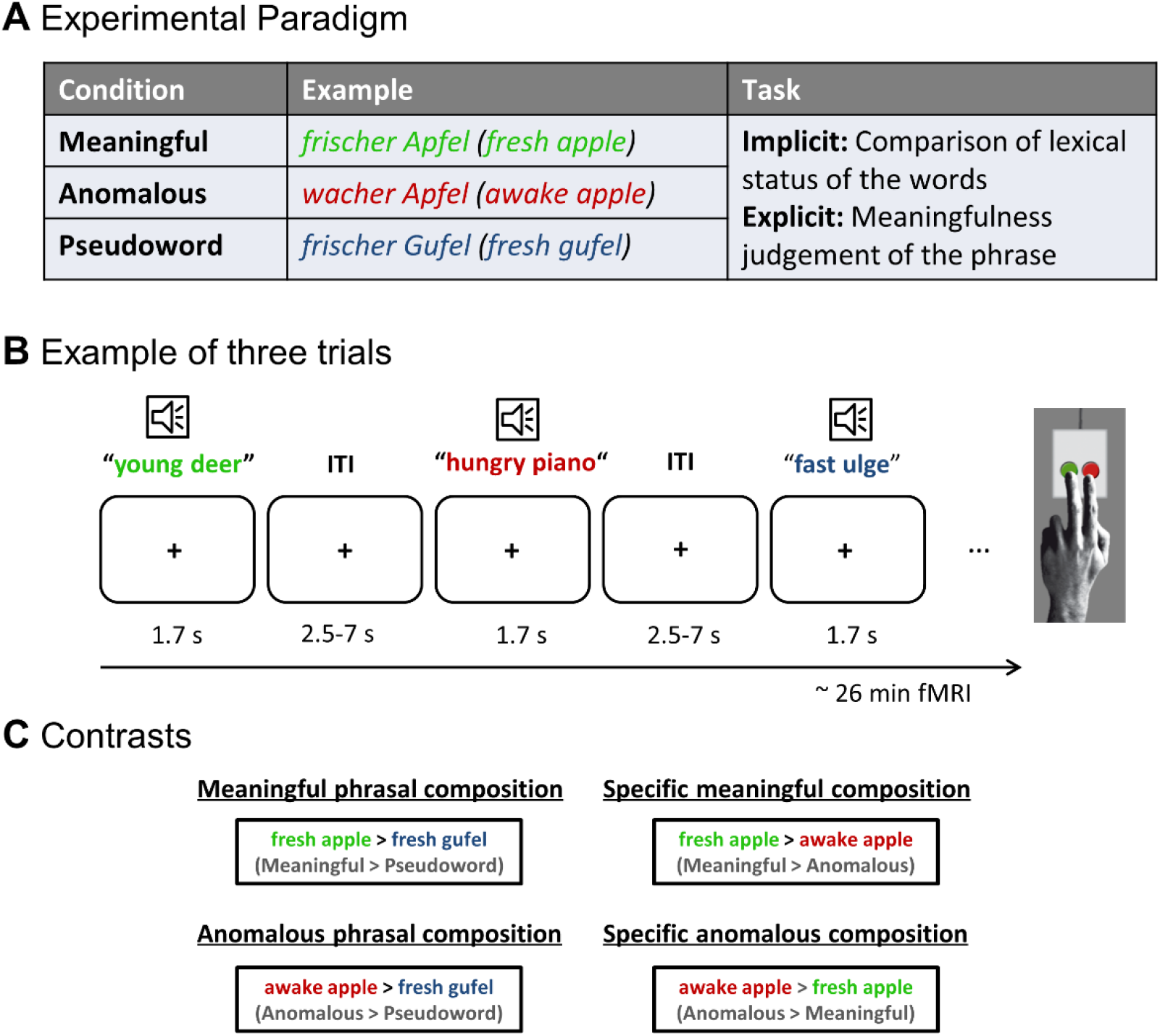
Experimental Design. (A) Experimental conditions and task descriptions used in the study. (B) Example of three trials. The inter-trial-interval (ITI) was jittered from 2500 ms up to 7000 ms with a mean duration of 4000 ms. (C) Contrasts of interest with involved compositional processes.

In the first session, participants performed an *implicit* task with respect to the meaning of the phrases. They were instructed to indicate whether both words had the *same* lexical status (i.e., both real words or both pseudowords) or whether they had a *different* lexical status (i.e., one real word and one pseudoword). Based on the vast literature on priming (cf. Lau et al., 2008), we expected our altered lexical decision task to capture automatic semantic processes. We added filler trials with two pseudowords and a pseudo-adjective paired with a real noun, to prevent participants from focusing on the second word only. In addition, single word trials served as control stimuli in order to balance the positive and negative responses. Participants were trained that single words required a “*different*” response, as there was no other word to compare it to. We note that this might not have been the intuitive response and made the task more difficult, however, we did not include the single word trials in any direct comparisons, as they served only as control trials.

In the second session, participants performed an *explicit* meaningfulness judgement task by indicating whether the phrase they heard was meaningful or not. To keep the amount of positive and negative responses equal, similarly to the implicit task, we additionally added single (meaningful) nouns as fillers which were not analyzed. Again, participants were trained to respond with “*meaningful*” to the single word condition. Before each session, participants completed a practice session outside of the scanner with a separate stimulus set. Subjects gave their response via button press of the left index or middle finger. Response button assignment was counterbalanced across participants. Both sessions consisted of 8 blocks with all conditions appearing 7 times in each block and pseudo-randomized with respect to order across participants. Blocks were separated by 20-second rest periods. Note that the implicit task was always performed in the first session, to keep the processing implicit and not biased by a previous task. The overall length of the two sessions was 31 minutes for the implicit task and 26 minutes for the explicit task (more filler trials in the implicit task, see *Stimulus* section).

### Stimuli

With our design we aimed to isolate the neural responses during the process of semantic composition, removing any effect of syntactic processing. To this end, we created a paradigm using one composition condition (*meaningful*, e.g. “fresh apple”) and two non-compositional conditions: In the *anomalous* condition, we created real word phrases that could not be combined using world knowledge (e.g. “awake apple”). In these stimuli, the adjective violated the selectional restriction criteria of the noun. The adjective “awake” typically describes living things but cannot be mapped onto non-living entities such as apples. In the *pseudoword* condition, we replaced the noun with a pseudonoun but kept the adjective the same. This way, the syntactic information is comparable for the pseudoword condition, but as it only contains one real word, a combination of concepts is not possible. This provided us with the advantage of avoiding confounds in the number of words presented and instead reduced the amount of conceptual information specifically.

To create real word stimulus pairs (*meaningful* and *anomalous*), we selected 400 nouns from the SUBTLEX-DE database (Brysbaert et al. 2011), constraining our search to the following criteria: mono- or disyllabic, masculine or neuter, monomorphemic, concrete, mean log frequency of 2.56. Concreteness was determined using an English corpus with ratings for 40.000 words (Brysbaert, Warriner, & Kuperman, 2014). As there is no existing large database for German, we translated the German corpus LANG (Kanske & Kotz, 2010) consisting of 1000 nouns into English and correlated the concreteness measures with each other. The correlation was very high between the two databases (r = 0.87, p < .001) and we thus used the English norms for our German words.

In the next step, we excluded all nouns that were ambiguous in their meaning. Orthographic neighborhood was controlled by calculating the Orthographic Levenshtein Distance 20 (OLD20, R package ‘vwr’), excluding all words deviating more than 1.5 times the interquartile range (IQR) from the mean OLD20 value across all items. Using the same parameters (frequency, number of syllables, OLD 20), except for the concreteness value specific to nouns, we selected adjectives that could modify concrete nouns. Even though we note that using adjectives which normally modify abstract nouns could have easily resulted in anomalous phrases, we decided against using those, to avoid having a confound of activation differences coming from the concreteness value.

To create meaningful word pairs, we generated all possible combinations of adjectives and nouns and assessed each pair’s frequency in the google web1t database (Linguistic Data Consortium, University of Pennsylvania). This database consists of n-gram counts of approximately 100 billion word tokens and searches for the frequency of each word pair. We excluded metaphoric pairs, alliterations and those that deviated more than 1.5*IQR from the mean pair frequency. We then created anomalous pairs by combining adjectives with nouns that did not occur in the google web1t output, taking care that each adjective combined with at least one noun meaningfully and anomalously and that the same held true for the nouns. In a pilot study, 20 participants who did not take part in the main experiment rated the plausibility of the remaining word pairs on a Likert scale from 1-6. We additionally asked for any potential associations to filter out items that could be understood metaphorically. From these ratings, we selected the highest rated pairs for the meaningful condition and the lowest rated pairs for the anomalous condition, excluding pairs that deviated more than 1.5*IQR from the mean of each condition. The final set of stimuli consisted of 56 phrases per condition. A list of psycholinguistic variables for both conditions can be found in Supplementary Table 1.

To create pseudowords that were comparable to real words, we used the pseudoword generator software Wuggy (Keuleers & Brysbaert, 2010). Here, we included all final real word stimuli and matched the pseudowords for length of subsyllabic segments, letter length, transition frequency and selected those items that deviated the least from the original words in their OLD20 value. Pseudo-homophones (i.e., stimuli that were pronounced as real words) were excluded.

Stimuli were recorded by a professional male speaker in a sound-attenuating chamber with a resolution of 16 bits and a sampling rate of 44.1 kHz. All words were spoken individually in the form of a statement. Thereafter, we cut all words into single files and normalized them to root-mean square amplitude using the Praat software (version 6.0.04). To keep the length of stimuli comparable, we concatenated all word-pairs with a constant noun onset at 1.1 seconds and a pause between the words of 40 ms, ensuring a natural sounding phrase. We furthermore included pseudoword-pseudoword and pseudoword-real-word pairs in the implicit task to ensure that participants could not make their judgment based on the second word only. Single nouns additionally served as fillers to keep the amount of positive and negative responses equal in both tasks. Fillers were not analyzed in the subsequent phases. The final set of stimuli had a mean length of 1703 ms (SD = 92 ms).

### fMRI acquisition

Functional images were acquired with a 3 Tesla Siemens Magnetom Scanner (Siemens, Erlangen) using a 32-channel head coil. To guarantee optimal signal of the ATL regions (Halai et al., 2014), we adopted a multiband dual gradient-echo echo planar imaging (EPI) sequence (60 slices in axial direction and interleaved order, TR = 2 seconds, short TE = 12 ms, long TE = 33 ms, flip angle of 90°, FOV = 204, slice thickness = 2.5 mm, interslice gap = 0.25 mm, multiband acceleration factor = 2) (Feinberg et al., 2010; Moeller et al., 2010). To further decrease artifacts in the ATL, slices were tilted by 10° off the AC-PC line. For offline distortion correction, field maps were acquired using a gradient dual-echo sequence (TR = 620 ms, TE1 = 4 ms and TE2 = 6.46 ms). Structural T1-weighted images were previously acquired and retrieved from the institute brain database for all participants. Images were acquired using an MPRAGE sequence (176 slices in sagittal orientation; TR: 2.3 s; TE: 2.98 ms; FoV: 256 mm; voxel size: 1 x 1 x 1 mm; no slice gap; flip angle: 9°; phase encoding direction: A/P).

### Behavioral Analysis

Analysis of the behavioral data was performed using the software R (Version 3.2.3). We calculated the mean percentage of correctly answered trials per participant and excluded any participant who performed with less than 75 % across all main conditions in any of the sessions (2 participants per session). For the analysis of reaction times, we only considered correctly answered trials within a response-time cutoff range of 2500 ms. All reaction times that deviated more than 3 SD from the mean per participant and condition were excluded (1.9 % in the explicit task, 1.8 % in the implicit task).

Statistical analyses were performed with the generalized linear mixed-effects model (GLMEM) using the lme4 package in R (Bates et al., 2014), assuming a Gamma distribution of our reaction time data. For the analysis of accuracy, we computed a mixed logit regression. We included by-participant intercepts to account for overall inter-individual differences and by-item intercepts and calculated two models with the respective reference levels *meaningful* and *anomalous*.

### fMRI Analysis

fMRI analyses were performed using SPM12 (Wellcome Trust Centre for Neuroimaging, http://www.fil.ion.ucl.ac.uk/spm/). The functional images from the two echoes were combined using a custom Matlab script that combined the images using a mean weighting by the temporal signal-to-noise ratio (tSNR) at each voxel. The combined functional images were then realigned to the first image, distortion corrected (using the field maps), co-registered to their corresponding structural image, normalized to MNI space (using a unified segmentation with a resampling size of 2.5 mm isotropic voxels) and smoothed with a 5 mm³ FWHM Gaussian kernel.

For statistical analyses, we estimated a general linear model (GLM) for each participant as implemented in SPM12, including one regressor for each condition and convolving the onset and duration of stimulus presentation with a canonical hemodynamic response function (HRF). Only correctly answered trials were analyzed and we added error trials as a regressor-of-no-interest. The 6 motion parameters were treated as nuisance regressors. A high-pass filter with 128 s cutoff was applied. For each subject we estimated the contrast for each condition against rest as well as direct contrasts between conditions.

At the group level, we conducted one-sample t-tests within each task (implicit and explicit) for each contrast. To identify brain regions that guide the successful composition of more complex meanings compared to simpler ones, we contrasted *meaningful* versus *pseudoword* phrases (*meaningful phrasal composition*; see Figure 1C). We also contrasted *anomalous* against *pseudoword* phrases to verify neural activity generated during the composition of complex meanings which do not result in meaningful phrases (*anomalous phrasal composition*). We further measured two more specific processes directly tackling semantic plausibility of complex meanings, by contrasting *meaningful* versus *anomalous* phrases (*specific meaningful composition*) and *anomalous* versus *meaningful* phrases (*specific anomalous composition*).

To explore regions that were activated independently of the final meaningfulness (*meaningful phrasal* and *anomalous phrasal composition*) in the explicit task, we performed a conjunction analysis based on the minimum statistic (Nichols et al., 2005), resulting in *general phrasal composition*. Furthermore, to detect regions that were activated independently of the task, we performed a conjunction analysis for the contrasts in both tasks.

Finally, to localize brain regions which responded significantly more to the explicit than the implicit task, we conducted paired t-tests. We therefore subtracted the contrast resulting from the implicit task from the one in the explicit, e.g., (meaningful_explicit_ > anomalous_explicit_) – (meaningful_implicit_ > anomalous_implicit_). These interactions were inclusively masked by the significant voxels of the minuend in order to restrict them to those voxels that were also activated in the task (cf. Hardwick, Caspers, Eickhoff, & Swinnen, 2018).

All contrasts were thresholded using a voxel-wise false discovery rate (FDR) correction of q < .05 with a cluster-extent threshold of 20 voxels to avoid reporting meaningless single voxel activations. Anatomical locations were identified using the SPM Anatomy Toolbox 2.2b (Eickhoff et al., 2005) and the Harvard-Oxford cortical structural atlas (https://fsl.fmrib.ox.ac.uk/fsl/fslwiki/).

### Psychophysiological interaction (PPI) analyses

Task-related functional connectivity between conditions of interest was assessed with a generalized psychophysiological interaction analysis (gPPI, McLaren, Ries, Xu, & Johnson 2012). Seed volumes of interest (VOI) were defined by drawing 6 mm spheres around each subject’s individual nearest activated voxel relative to the group peak of a given contrast at a threshold of p < 0.05. This lenient threshold ensured that each participant’s VOI was in the same anatomical region as the group peak. To explore functional connectivity between key semantic regions and other brain areas we seeded from the following activation peaks: left aIFG, PGa, PGp, pMTG, ATL and DMPFC. The design matrix of each participant for each VOI comprised (1) the deconvolved time series of the first eigenvariate of the BOLD signal from the VOI, forming the physiological variable, (2) each condition convolved with the HRF, forming the psychological variable, and (3) the interaction of the psychological and physiological variable, forming the PPI term. At the single-subject level, whole-brain GLMs were conducted creating 3 contrasts (of the PPI terms) for each VOI model based on the univariate GLM results: 1) meaningful > anomalous, 2) meaningful > pseudowords, 3) anomalous > pseudowords. At the group level, we conducted one-sample t-tests for each contrast of interest. Contrast images were thresholded at p < 0.05, cluster-level family wise error (FWE) corrected, with a voxel-wise threshold of p < 0.001.

## Results

### Behavioral results

Overall accuracy was high in both tasks (mean implicit: 94.14%, mean explicit: 95.75%). In the implicit task (comparison of lexical status of the words), meaningful phrases had a significantly higher accuracy than both anomalous and pseudoword phrases. In the explicit task (meaningfulness judgement of the phrase), an opposite picture emerged with pseudoword phrases being significantly more accurate than anomalous and meaningful phrases (Figure 2A/C; SI Table 2). Reaction times in the implicit task were significantly faster for meaningful than anomalous and pseudoword phrases and for anomalous relative to pseudoword phrases. In the explicit task, reaction times were significantly faster for pseudoword phrases than meaningful and anomalous and for meaningful than anomalous phrases (Figure 2B/D; SI Table 2).

**Figure 2.**
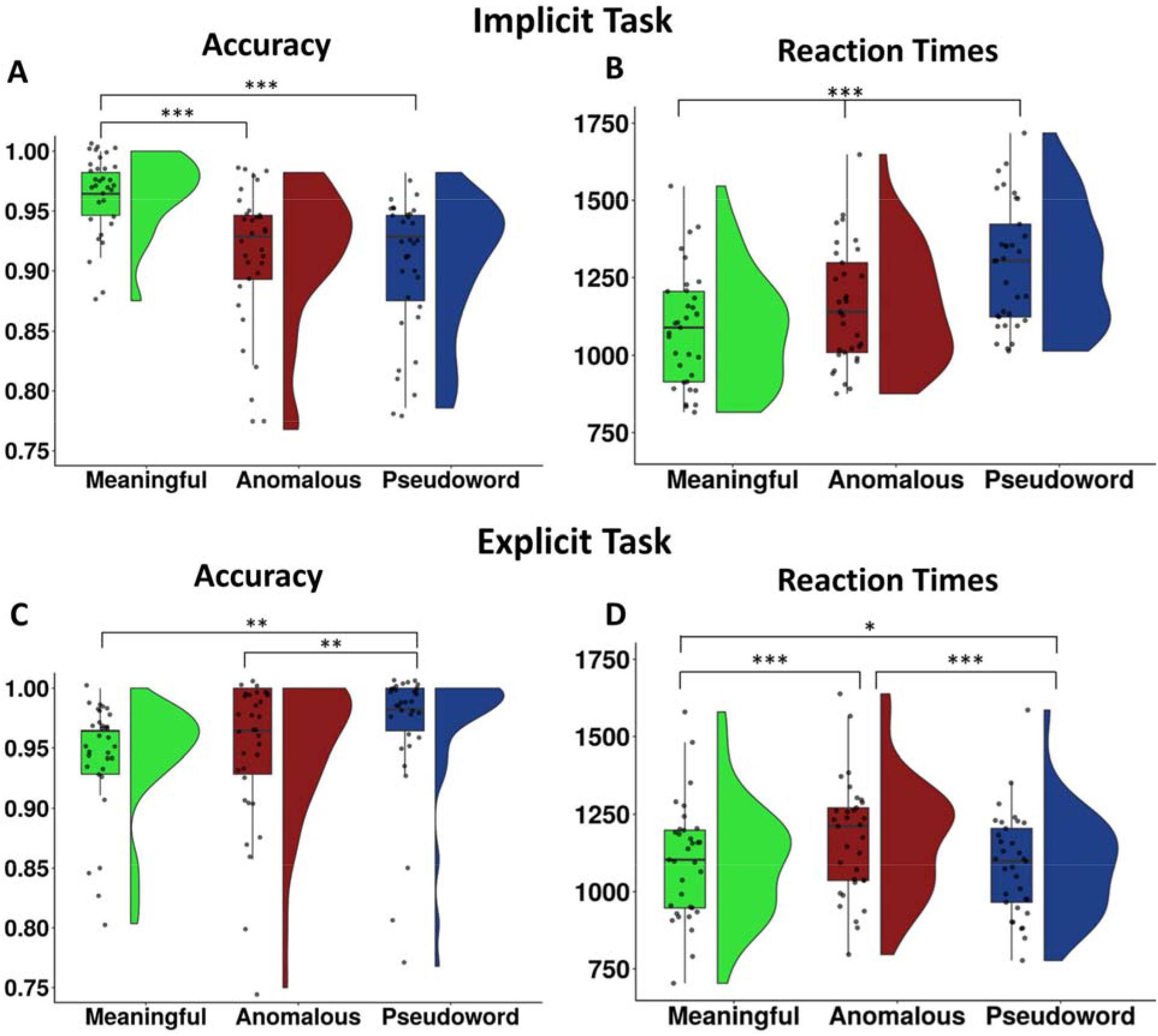
Raincloud plots. (Allen et al., 2019) illustrating the data distribution of each participant’s mean behavioral scores and boxplots overlaid with individual mean data points for the implicit (A, B) and explicit task (C, D). * *p* < .0.05; ** *p* < 0.01; *** *p* < 0.001.

### fMRI results

#### Implicit Task: Meaningful phrasal composition

In the implicit task, only the contrast of meaningful > pseudoword phrases (*meaningful phrasal composition*) yielded significant results. Here, we found increased activation in left AG (PGp), dorsomedial prefrontal cortex (DMPFC), pMTG/ITG and a small cluster in right anterior cingulate cortex (ACC) (Figure 3; SI Table 3).

**Figure 3.**
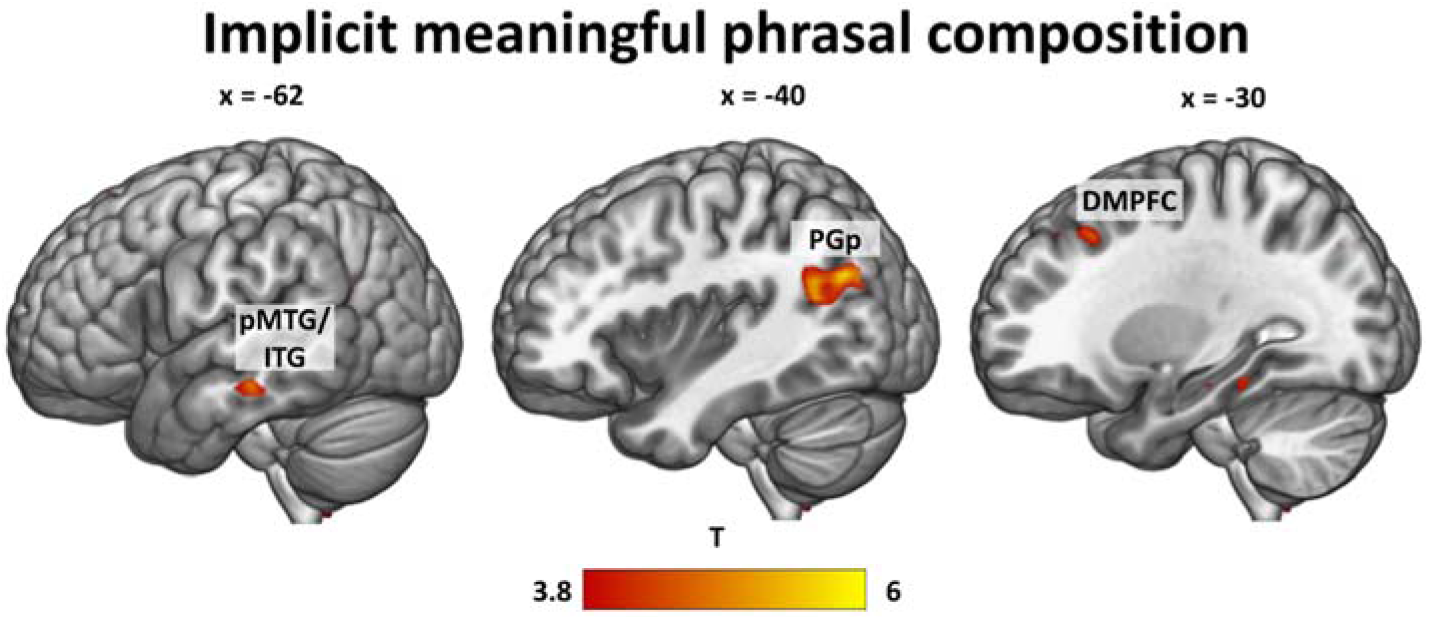
**Whole-brain activations in the implicit task** for the contrast meaningful > pseudoword phrases, thresholded at q < 0.05 FDR-corrected. DMPFC = dorsomedial prefrontal cortex, pMTG = posterior middle temporal gyrus, pITG = posterior inferior temporal gyrus, PGp = angular gyrus posterior division.

#### Explicit Task: Meaningful phrasal composition

Comparing meaningful phrases with pseudoword phrases in the explicit task yielded activation in a wide-spread largely left lateralized network of regions comprising aIFG (pars orbitalis), DMPFC, AG (PGp) extending into PGa, SMG and IPS, pMTG, pITG, ATL (including temporal pole), ACC (extending to right ACC) and posterior cingulate cortex (PCC), cerebellum (crus I/II), precuneus, insula and hippocampus. Additionally, increased right-hemispheric activity was found in the cerebellum, insula (extending into temporal pole), primary motor area (M1), fusiform gyrus, pITG and AG (PGp) (Figure 4A; SI Table 4).

**Figure 4.**
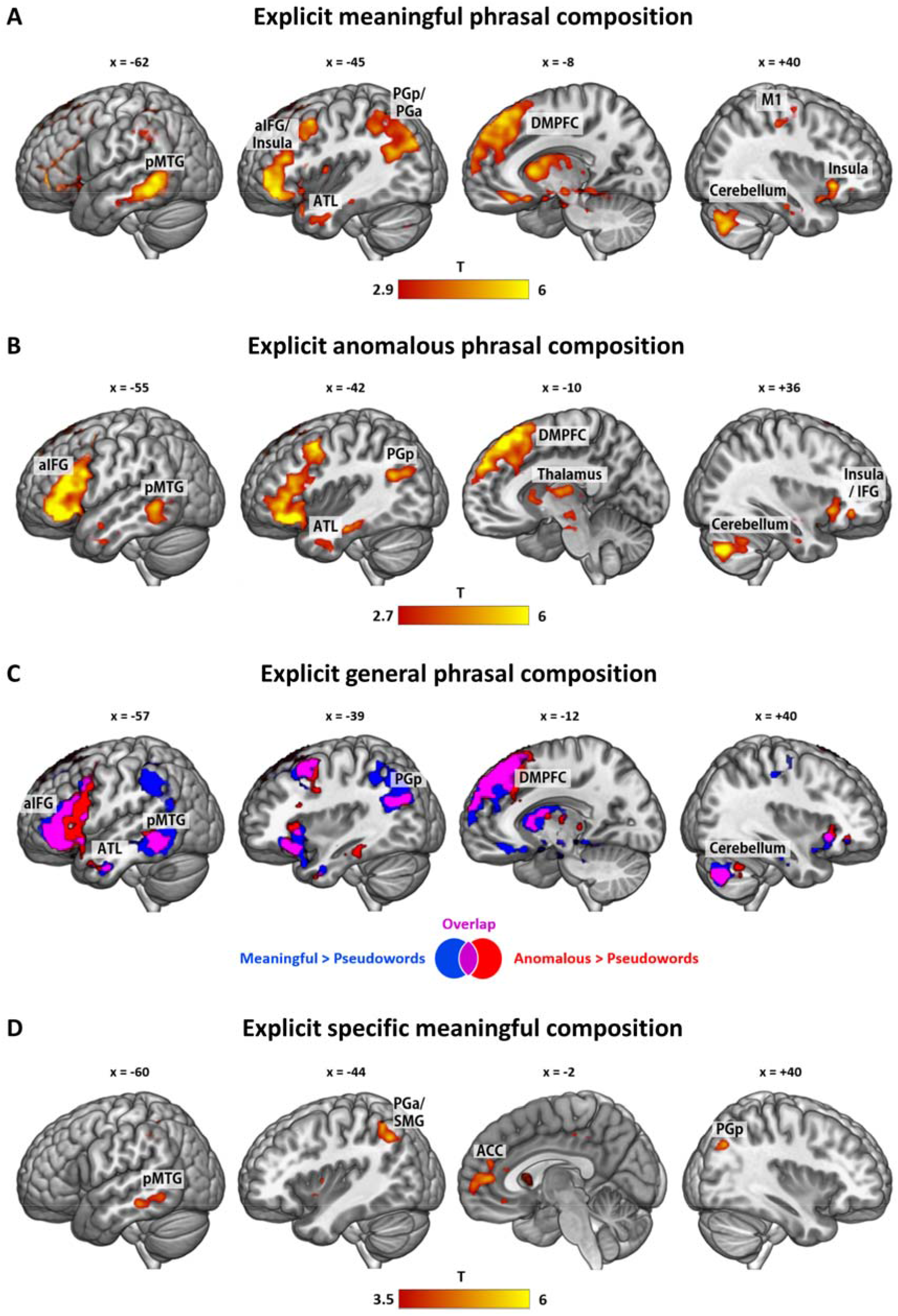
**Whole-brain activations in the explicit task** for the contrast (A) meaningful > pseudowords, (B) anomalous > pseudowords, (C) Overlap (purple) of the contrasts meaningful > pseudowords (blue) and anomalous > pseudowords (red), resulting in *general phrasal composition* and (D) the specific contrast of meaningful > anomalous phrases. All activation maps are thresholded at q < 0.05 FDR-corrected. ACC = anterior cingulate cortex, aIFG = anterior inferior frontal gyrus, ATL = anterior temporal lobe, DMPFC = dorsomedial prefrontal cortex, pMTG = posterior middle temporal gyrus, PGa = angular gyrus anterior division, PGp = angular gyrus posterior division, SMG = supramarginal gyrus.

#### Explicit Task: Anomalous phrasal composition

A similar pattern emerged for the contrast of anomalous versus pseudoword phrases. This contrast yielded a largely left lateralized network of regions including aIFG (pars orbitalis/triangularis), DMPFC, pMTG, ATL, AG (PGp), fusiform gyrus and thalamus. Right hemispheric activation comprised the cerebellum, aIFG/insula, amygdala and thalamus (Figure 4B; SI Table 5).

#### Conjunction analysis: explicit meaningful phrasal composition and anomalous phrasal composition (general phrasal composition)

To explore regions that are activated independently of the meaningfulness of the final phrase, we conducted a conjunction analysis of the contrasts meaningful > pseudowords ∩ anomalous > pseudowords. This conjunction will be referred to as *general phrasal composition*, to emphasize that it does not depend on the meaningfulness. This yielded common activations for all regions that were also involved in anomalous > pseudoword processing, showing that *anomalous phrasal composition* engages a subset of the regions for *meaningful phrasal composition* (Figure 4C; SI Table 6). Thus, the left aIFG, ATL, pMTG, PGp, DMPFC, thalamus, right aIFG and cerebellum appear to be involved in phrasal composition independently of the plausibility of the resulting phrase.

#### Explicit Task: Specific Meaningful Composition

To determine regions that guide *specific meaningful composition*, we compared meaningful versus anomalous phrases in the explicit task. Here, we found significantly increased activity in the anterior part of left AG (PGa) extending into supramarginal gyrus (SMG) and intraparietal sulcus (IPS), ACC, left pMTG, left ventromedial prefrontal cortex (vmPFC) and a small cluster in the posterior part of the right AG (PGp) (Figure 4D; SI Table 7).

The opposite contrast of anomalous versus meaningful phrases did not yield any significant activation differences.

#### Task-independent activation for meaningful phrasal combination

After exploring the contrasts for each task separately, we were interested in common regions that are activated independently of task. To this end, we conducted conjunction analyses of the only contrast that yielded significant results in both tasks: explicit *meaningful phrasal composition* ∩ implicit *meaningful phrasal composition.* We observed a significant cluster in the left PGp and very small clusters in the left pITG and DMPFC (Figure 5; SI Table 8). Thus, the only regions showing task-independent activations for *meaningful phrasal composition* are the posterior part of the AG and to a lesser extent, parts of DMPFC and pITG.

**Figure 5.**
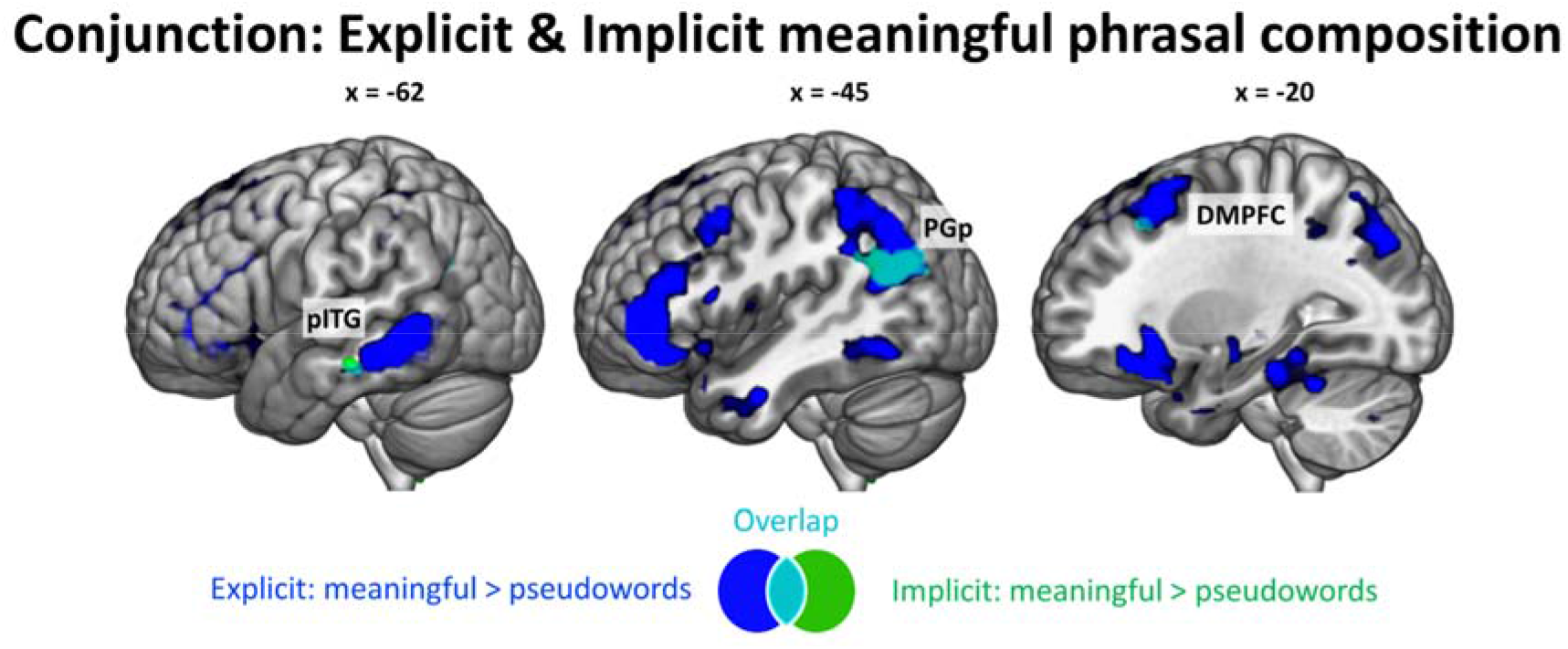
Task independent activations. Overlap (cyan) between the contrast meaningful > pseudowords in the explicit task (blue) and the implicit task (green), thresholded at q < 0.05 FDR-corrected. DMPFC = dorsomedial prefrontal cortex, pITG = posterior inferior temporal gyrus, PGp = angular gyrus posterior division.

#### Task-specific activations

From the results above, it appears that brain regions exist which are selectively involved during the explicit task, but not during the implicit one. To further substantiate this finding, we conducted paired t-tests for all contrasts between the two tasks.

For the contrast of *meaningful phrasal composition*, paired t-tests confirmed that left aIFG (extending into insula), DMPFC, pMTG/ITG, IPS/SMG and right aIFG, thalamus and cerebellum are significantly more involved in the explicit than in the implicit task (Figure 6A; SI Table 9).

**Figure 6.**
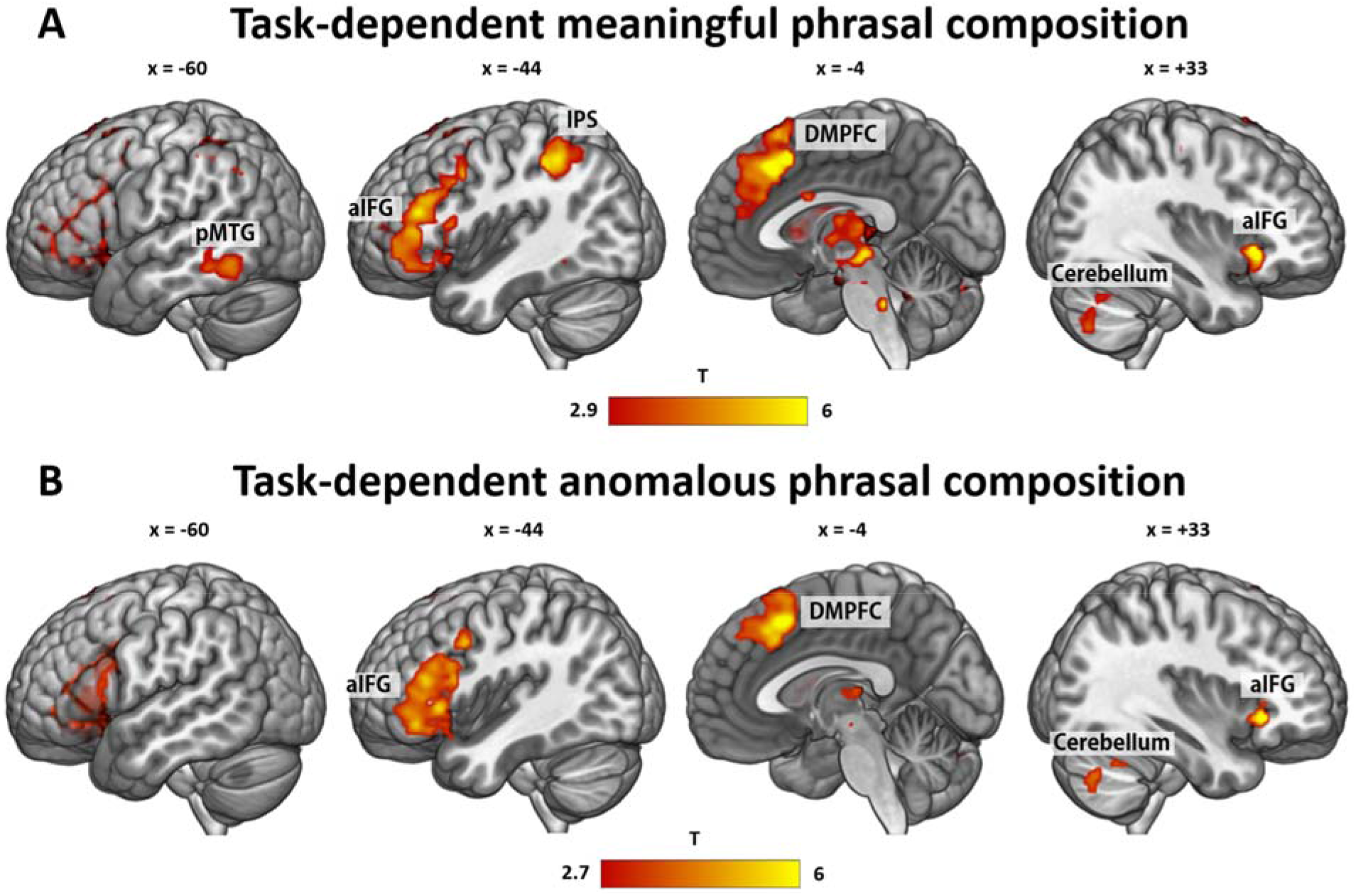
Task-dependent activations. (A) Stronger activation during explicit than implicit task for meaningful > pseudowords phrases. (B) Stronger activation during explicit than implicit task for anomalous > pseudoword phrases. Thresholded at q < 0.05 FDR-corrected. aIFG = anterior inferior frontal gyrus, DMPFC = dorsomedial prefrontal cortex, IPS = inferior parietal sulcus, pMTG = posterior middle temporal gyrus.

*Anomalous phrasal composition* also revealed task-dependent activations in left aIFG, DMPFC, right cerebellum, aIFG and thalamus (Figure 6B, SI Table 10). Thus, these regions seem to be selectively involved when explicit meaningfulness judgement is required. For *specific meaningful composition*, this only yielded significant activations when lowering the threshold to cluster-level FWE correction (p < 0.05) in PGa and ACC. Thus, we can only cautiously speak of a trend of task dependence in these regions.

#### Psychophysiological interaction (PPI) analysis for meaningful phrasal composition

Finally, we set up several PPI models to investigate task-specific interactions. In the explicit task, we found significant functional coupling between the left PGp (as seed region) and left aIFG (pars triangularis; BA45) and bilateral pre-supplementary motor cortex for *meaningful phrasal composition* (i.e. meaningful > pseudowords; Figure 7; SI Table 11). No other seed region or contrast yielded significant results.

**Figure 7.**
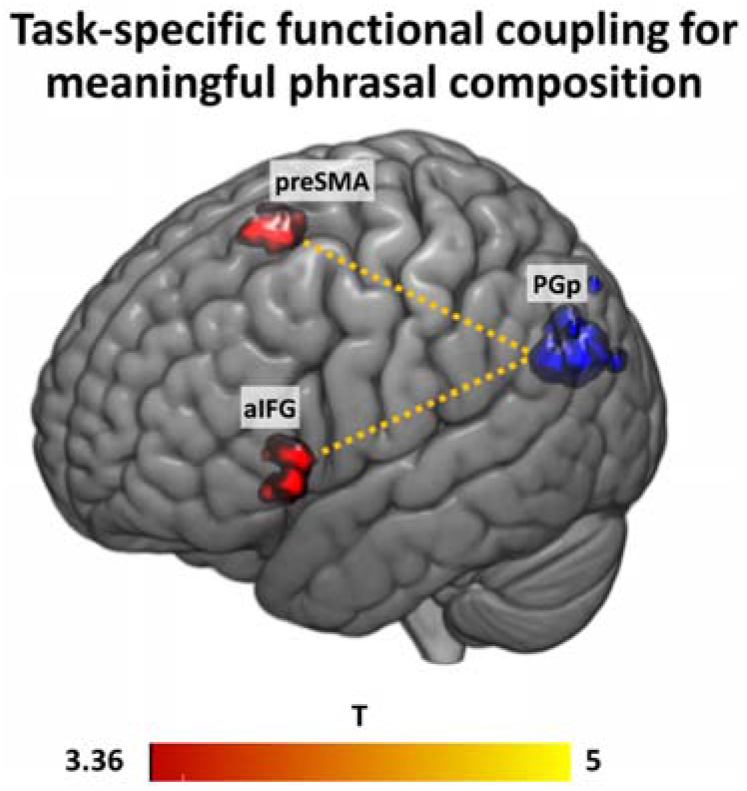
gPPI results with PGp as seed region (blue; 6 mm spheres around individual peak activation overlaid on top of each other) for the contrast meaningful > pseudowords in the explicit task (thresholded at p < 0.05 FWE-corrected at the cluster level).

## Discussion

In the present study, we sought to characterize the neural correlates of basic semantic composition and the task-dependent interaction network characterizing such process. Despite numerous efforts to identify the key regions of combinatorial semantic processing in previous neuroimaging studies, several questions remained open.

Here, we developed a paradigm that is sensitive to capture both the general process of combining two real words together, comprising *meaningful phrasal-* and *anomalous phrasal composition*, as well as the more specific process of successful combination of two words into a meaningful phrase (*specific meaningful composition*). We identified separable neural correlates for these two processes. The more *general phrasal composition* process, which appears to be largely independent of the plausibility of the resulting phrase, is associated with a widely distributed pattern of left-lateralized activity, including the aIFG, ATL, DMPFC, pMTG and AG. Crucially, the posterior part of the AG is involved in a task-independent manner, suggesting a role in general semantic representation processes that goes beyond task-specific activation. This region might thus reflect automatic general phrasal composition. In contrast, aIFG, a large part of DMPFC, pMTG and IPS/SMG show stronger engagement for explicit *meaningful phrasal composition*, than in the supposedly more lexical-level implicit task. *Meaningful phrasal composition* furthermore relies on the functional interaction between the left posterior AG, the aIFG and the pre-supplementary motor area, as shown in our PPI analysis. In contrast, *specific meaningful composition* engages a subset of regions from the phrasal composition pattern (anterior AG, pMTG, ACC). These regions thus seem to lie at the heart of successful meaning composition, necessary for evaluating the plausibility of a phrase.

While previous research has proposed aIFG, ATL and AG to play a key role in semantic composition, our results help to disentangle the subprocesses that might guide successful semantic composition in these regions. Comparing the two real-word conditions (i.e., meaningful and anomalous phrases) to the pseudoword condition (referred to as *general phrasal composition* in our study) requires the successful retrieval of two separate concepts compared to just one concept, thus a general lexical-semantic retrieval process appears to be involved. Conversely, *specific meaningful composition* reflects the “pure” process of conceptual combination. Since this process can be regarded as the more specific form of phrasal composition, it is not surprising that we only find a subset of regions from the *meaningful phrasal composition* contrast involved, including the anterior AG.

### General phrasal composition engages a large semantic network in the left hemisphere

In line with previous research, we found recruitment of left AG, aIFG and ATL, along with other classical regions of the semantic network (e.g., left pMTG and DMPFC) and right-hemispheric motor (control) regions (cerebellum, premotor, primary and somatosensory motor cortex) for the contrast of meaningful versus pseudoword phrases (i.e., *meaningful phrasal composition*) in the explicit task. Additionally, conjunction analyses revealed that the activation pattern largely overlapped with the contrast of anomalous > pseudoword phrases (i.e., *anomalous phrasal composition*), showing that these regions are involved in *general phrasal composition*, independently of the plausibility of the resulting phrase.

Consistent with its proposed role in executive semantic control (Chiou et al., 2018; Lambon Ralph et al., 2016; Noonan et al., 2013; Whitney et al., 2011), the left aIFG was selectively activated in the explicit task, for both real-word conditions compared to the pseudoword condition. Previous studies have associated the left aIFG with monitoring and selecting semantic information when several alternatives are present (Binder & Desai, 2011; Lau et al., 2008; Noonan et al., 2013; Whitney et al., 2012). In our study, the aIFG activates as a function of the amount of words that can be retrieved from the lexicon, independent of the meaningfulness of the final phrase as it was observed both during *meaningful phrasal-* and *anomalous phrasal composition*. This fits with the results from Schell et al. (2017), who found aIFG activation for adjective-noun phrases (*blue boat*) versus single words (*boat*). Consequently, aIFG involvement might reflect increased semantic load carried by real words, which also goes beyond sentential level (Zaccarella et al., 2015), to track the amount of semantic information to be integrated.

Unlike previous studies, which have suggested the ATL to be the key conceptual composition hub (e.g. Baron et al. 2011; Bemis & Pylkkänen 2011; Westerlund & Pylkkänen 2017), we found evidence for its contribution only at the more general phrasal level in the explicit task (i.e., for the contrasts meaningful > pseudowords and anomalous > pseudowords) and not for *specific meaningful composition*. Based on our findings, we infer that the ATL may code for the amount of conceptual information that can be retrieved rather than guiding the conceptual combination of two concepts into a whole. Furthermore, our results do not lend support to the notion that the ATL guides the automatic composition of concepts (Bemis & Pylkkänen, 2013a), as we only find involvement of this region in the explicit task.

Another region in the temporal lobe that was selectively activated during the explicit task is the posterior MTG, reaching into ITG. As it was also engaged during *specific meaningful composition*, we will turn to its discussion below.

Our study additionally revealed a strong engagement of DMPFC during explicit *general phrasal composition*. In their meta-analysis, Binder and colleagues (2009) defined the DMPFC as one of the core semantic regions often activated by semantic tasks. However, previous reviews have failed to acknowledge its role in semantic tasks and a consistent theory of its exact contribution to semantic processing is still lacking. Binder and Desai (2011) proposed that the DMPFC, together with the left IFG, acts as control region that guides goal-directed retrieval of conceptual information stored in temporo-parietal cortices. Additional evidence comes from Graves et al. (2010), who found DMPFC engagement for meaningful as compared to reversed noun-noun phrases only in their explicit task. Consequently, DMPFC seems to be a semantic control region that has been overlooked previously but should be granted a more central role in semantic processing, including semantic composition.

Regarding the role of different AG subregions in semantic processing, previous meta-analyses have identified a dorsal, middle and ventral subdivision, each serving a different function with respect to semantic tasks (Noonan et al., 2013; Seghier, 2013). Crucially, our activation cluster for *general phrasal composition* in the posterior division PGp overlaps with the mid-AG region identified in the meta-analysis by Noonan et al. (2013) (MNI coordinates x, y, z: −39, −69, 30) and Seghier et al. (2010) (MNI coordinates x, y, z: −48, −68, 29). Despite partly contradictory findings, this subregion was consistently reported for processing concrete relative to abstract concepts and was thus ascribed a role in semantic representation for rich multimodal concepts. As we found relatively stronger involvement of PGp in both real word conditions as compared to the pseudoword condition, our results fit with the account that this subregion codes for the semantic richness. More conceptual information can be retrieved during the two real-word conditions than for the pseudoword condition, but this distinction does not hold when the two real word conditions are compared directly. Thus, we did not observe engagement of left PGp in *specific meaningful composition* but only in *general phrasal composition*. We believe that our results help to specify the role of different subregions in the left AG in semantic composition. The posterior cluster seems to guide the more general semantic integration of two concepts, while the anterior part evaluates the meaningfulness of the resulting phrase. Note further that PGp, together with small clusters in the pITG and DMPFC, were the only regions that were also involved in the implicit task, speaking for a task-independence of these regions in lexical-semantic processes. Thus, in our view, PGp is involved in the representation of more versus less conceptual information regardless of the plausibility of the resulting phrase and independent of the task. Conversely, a region in the IPL that shows strong task-dependent involvement is the IPS, bordering posterior SMG. This region was significantly more involved during the explicit than the implicit task and thus likely guides controlled conceptual retrieval together with the aIFG and DMPFC.

### Fronto-parietal interactions during meaningful phrasal composition

Aside from task-related activations, our results revealed that during the processing of meaningful phrases, PGp shows increased functional connectivity with aIFG and pre-SMA. The task-related functional coupling was stronger for the processing of meaningful phrases compared to pseudoword phrases. These findings extend previous results of functional connectivity at the single word level. In a TMS study, Hartwigsen et al. (2015) showed that temporary disruption of either aIFG or AG alone did not lead to a significant impairment of semantic decisions, while combined TMS over both regions significantly delayed reaction times in the semantic task. This suggests that AG and aIFG can compensate for the disruption of the respective other node. Moreover, after longer-lasting disruption of left AG, this area had an inhibitory influence on the left aIFG during semantic word decisions, further substantiating the strong interaction between both regions (Hartwigsen et al., 2017). Crucially, our results provide novel supporting evidence for the notion of a semantic network involving AG and aIFG not only at the single word, but also at the basic combinatorial level. Aside from aIFG, task-related functional coupling was also increased between PGp and the pre-SMA. While the pre-SMA does not belong to the classical language network, there is evidence that it is involved in higher-order cognitive processes such as semantic processing, independent of motor effects (Hertrich et al., 2016). In summary, the results from our functional connectivity analysis significantly extend previous findings of changes in task-related activity during semantic composition and provide new insight into functional interactions during meaningful phrasal compositions at the network level. We note that one limitation of our PPI analyses is that from the 6 seed regions, only the PGp yielded significant connectivity with any other brain region.

### Specific meaningful composition engages the anterior AG and semantic control regions

When directly comparing meaningful phrases with anomalous phrases, we found increased activity in several semantic regions, mainly in the left hemisphere. These regions included the left anterior AG (PGa) and neighboring SMG / IPS, the left pMTG, the bilateral ACC and the left vmPFC, as well as several smaller clusters in the right hemisphere, including the right posterior AG.

The strong contribution of the anterior part of the angular gyrus (PGa) is well in line with a previous study that used a similar task (Price et al., 2015). In that study, AG (with peak in PGa) showed increased activity for more meaningful relative to less meaningful adjective-noun phrases. Recently, left AG was found for the processing of verb phrases and noun phrases relative to word lists (Matchin, Liao, et al., 2019), providing further evidence for its key role in successful conceptual composition. The functional relevance of the left AG for semantic composition was demonstrated in lesion and neurostimulation studies (Price et al., 2015, 2016). Crucially, our activation cluster for specific meaningful composition was located at the border of PGa, posterior SMG (PFm) and IPS. This overlaps with the dorsal AG subregion identified by Noonan et al. (2013) that was associated with semantic control and conceptual combination. PGa thus seems to guide explicit meaningful composition that leads to a successful new concept by evaluating the plausibility of the phrase. Aside from the left AG, our results also revealed a small cluster in the right AG during *specific meaningful composition*. This activation is in line with two previous studies that reported right AG involvement in basic combinatorial processing (Graves et al. 2010; Price et al. 2015). Graves and colleagues proposed that while left temporo-parietal regions represent single word meaning, the right hemisphere represents the overlap of single concepts and combines the meaning of them, which might explain the observed upregulation of the right AG in the present study.

Another semantic region that we found for *specific meaningful composition* was the left pMTG. This finding was a bit surprising, as the pMTG has not classically been ascribed a key role in semantic composition. Rather, converging evidence from neuroimaging and neurostimulation studies has attributed the pMTG (together with the anterior IFG) a role in semantic control during semantic association tasks at the word level (Davey et al., 2016; Noonan et al., 2013; Whitney et al., 2011). Our results fit with this view insofar as we found much stronger activation in pMTG in the explicit than the implicit task. It is reasonable to assume that the implicit task does not require the same level of executive semantic processing as the explicit task. The observed engagement of the pMTG in all real-word contrasts (i.e., in explicit *specific meaningful composition*, *meaningful phrasal composition* and *anomalous phrasal composition*) might thus indicate that this region generally guides the successful retrieval of concepts.

Interestingly, we also found engagement of bilateral ACC during *specific meaningful composition*. This region is typically involved in error monitoring tasks (Botvinick et al., 2004), a process that is unlikely to occur during meaningful composition. It is furthermore a key region in the cingulo-opercular control network and associated with task maintenance (Vaden et al., 2013). However, there is recent evidence that ACC is also involved in semantic tasks. Almeida and colleagues (2016) found widespread activations including ACC for indeterminate sentences (*The author began the book*) as compared to preferred sentences (*The author wrote the book*) and anomalous sentences (*The author drank the book*). They interpret the role of these regions as employing pragmatic-inferential processes. Furthermore, the meta-analysis by Noonan et al. (2013) identified the ACC as part of the wide-spread semantic control network. The fact that we found ACC involvement only in the explicit task supports this view. Finally, we also observed a small cluster in the left vmPFC for the contrast of *specific meaningful composition*. This region, has been found in a number of MEG studies and a recent review by Pylkkänen (2019), attributes the vmPFC the role of representing the final combinatory output.

Notably, we did not find significant task-related activity during implicit *specific meaningful composition*. The lack of activation differences in the implicit task could initially suggest that meaning is not automatically composed but might rather be restricted to situations where it is task-relevant. However, the observed significant differences in accuracy and response times between meaningful and anomalous phrases in the implicit task might indicate that participants did automatically evaluate the meaning of the phrases. Crucially, the lexical material of meaningful and anomalous phrases did not differ, so participants should be equally successful in deciding whether the stimuli are both real words or not. The observed higher error rate for anomalous phrases thus points towards an automatic and implicit meaningfulness judgement. A possible explanation for why we did not find activation differences in the fMRI results could be that the nature of our lexical status task required increased task demands, which was overall similar for meaningful and anomalous phrases. An intriguing question for future studies would be whether other implicit semantic tasks (e.g. classical lexical decision, phonological tasks) are able to detect combinatorial processing differences in fMRI data for meaningful versus anomalous phrases, which, in the current study, are only visible at the behavioral level.

Finally, it should be noted that we did not observe significant activation differences for the reversed contrast in the explicit task (i.e., for *anomalous* relative to *meaningful* phrases). While it is conceivable to find a similar effect as reported for the classical N400 in the electroencephalography (EEG) literature, previous fMRI studies have also found that the effect is a lot weaker in the hemodynamic modality (Lau & Namyst, 2019). This could, among other reasons, be due to the much lower time-sensitivity of fMRI compared to EEG, with the latter thus being better suited to capture the relatively short-lasting N400 effect.

Another possible explanation is the nature of our task: Having to judge the meaningfulness might imply that there would be meaning in the phrases. Participants actively searched for associations and meaning to make their judgement and were thus already primed towards the more meaningful condition (Kuperberg, 2007).

## Conclusion

In the present study, we identified distinct neural signatures for two processes during explicit basic semantic composition: a *general phrasal composition* process, which is independent of the plausibility of the resulting phrase, and a *specific meaningful composition* process, strongly dependent on the resulting plausibility. *General phrasal composition* engages a widespread semantic network in the left hemisphere, including the posterior AG (PGp), the aIFG, DMPFC and large parts of the temporal lobe. Crucially, only the PGp shows task-independent engagement, pointing towards a role in automatic semantic processing. PGp furthermore strongly interacts with pre-SMA and another key semantic region in the left hemisphere, the aIFG, thus forming the core semantic network of successful meaningful phrasal composition. The more *specific meaningful composition* engages a subset of the semantic control regions found for phrasal composition, and the left anterior AG (PGa). Consequently, the AG appears to be decomposable into distinct subregions during semantic composition: PGp codes for the amount of conceptual information present in the phrase, while PGa successfully combines the meaning of two separate concepts to a whole.

## Acknowledgments

We wish to thank Lisa Kunz for help during data acquisition and Nicole Pampus for acquiring participants. We also like to thank Laura Nieberlein for her assistance in piloting the experiment. Moreover, we thank Angela D. Friederici for helpful discussions and Philipp Kuhnke and Anna Rysop for comments on this manuscript. This work was supported by the Max Planck Society. GH is supported by the German Research Foundation (DFG, HA 6314/3-1, HA 6314/4-1). We thank the University of Minnesota Center for Magnetic Resonance Research for the provision of the multiband EPI sequence software.

