## Supplementary Materials for "Differential contributions of left-hemispheric language regions to basic semantic composition"

**Supplementary Material**

**Supplementary Materials and Methods**

**Supplementary Table 1.** Psycholinguistic variables for the two real-word conditions.

|  | Anomalous | Meaningful | *p* |
| --- | --- | --- | --- |
| **Adjectives** | | | |
| Frequency | 2.13 (0.96) | 2.11 (0.95) | 0.900 |
| OLD-20 | 1.98 (0.45) | 1.91 (0.43) | 0.400 |
| **Nouns** | | | |
| Frequency | 2.57 (0.59) | 2.57 (0.55) | 0.960 |
| OLD-20 | 1.66 (0.35) | 1.67 (0.31) | 0.830 |
| Concreteness | 4.86 (0.13) | 4.85 (0.15) | 0.720 |
| **Pairs** |  |  |  |
| Meaningfulness rating | 1.47 (0.31) | 5.49 (0.36) | **< 0.0001** |
|  | *1.34 (0.24)* | *5.69 (0.27)* |  |

Frequency and OLD-20 (orthographic neighborhood) measures were taken from the SUBTLEX-DE database and frequency is given as log-transformed per 1 million words. Concreteness was determined using concreteness ratings for 40.000 English words. Ratings were obtained from 20 participants who did not take part in the fMRI experiment. Ratings in italics are averaged ratings of the fMRI participants in a post-hoc questionnaire. They indicate great overlap with our predefined conditions.

**Supplementary Results**

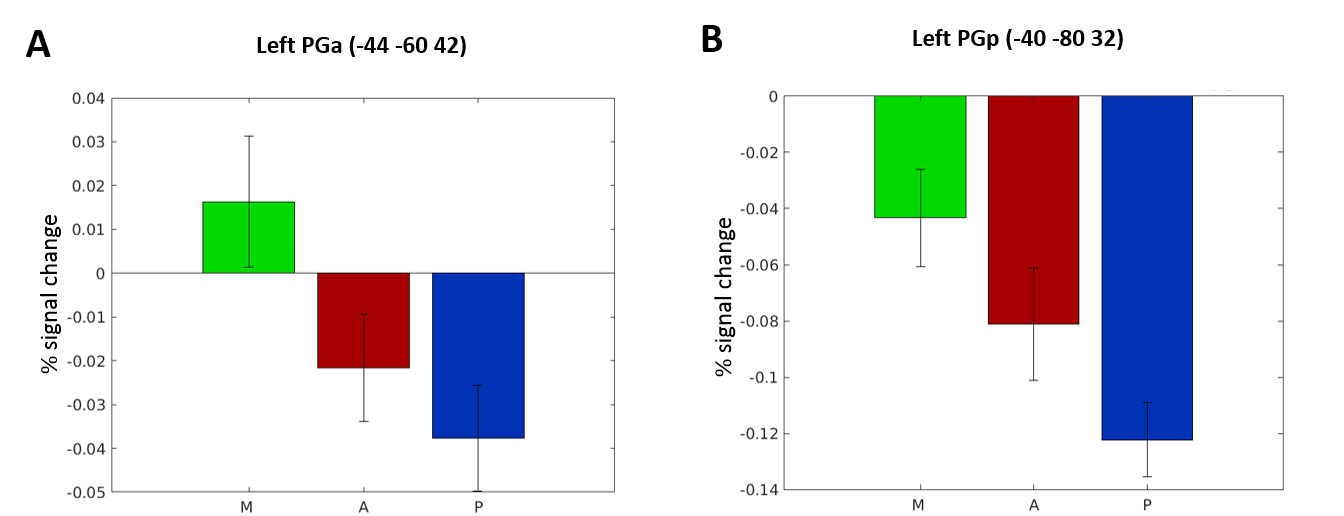

**Supplementary Figure 1**. Percent signal change for PGa (A) and PGp (B) for each condition compared to the implicit rest baseline. 6-mm spheres centered on the peak voxel from the contrast meaningful > anomalous for PGa and from meaningful > pseudowords for PGp were created using the MarsBaR toolbox (version 0.44; <http://marsbar.sourceforge.net/>).

**Supplementary Table 2.** Behavioral results.

|  | | **IMPLICIT** | | | | | **EXPLICIT** | | | |
| --- | --- | --- | --- | --- | --- | --- | --- | --- | --- | --- |
| **Predictor** | *Coef. ß* | | *SE(ß)* | *T* | *p* | *Coef. ß* | | *SE(ß)* | *T* | *p* |
| **Accuracy** |  | |  |  |  |  | |  |  |  |
| m > a | 1.02 | | 0.23 | 4.42 | **< 0.001** | -0.09 | | 0.18 | -0.49 | 0.62 |
| m > p | 1.14 | | 0.23 | 4.97 | **< 0.001** | -0.59 | | 0.19 | -3.08 | **< 0.01** |
| a > p | 0.12 | | 0.2 | 0.59 | 0.556 | -0.51 | | 0.19 | -2.6 | **< 0.01** |
| **Reaction Time** | | | | | | | | | | |
| m > a | 76.32 | | 4.28 | 17.84 | **< 0.001** | 67.99 | | 3.82 | 17.8 | **< 0.001** |
| m > p | 204.44 | | 5.14 | 39.76 | **< 0.001** | -8.59 | | 3.6 | -2.39 | **< 0.05** |
| a > p | 128.12 | | 6.53 | 19.61 | **< 0.001** | -76.58 | | 4.23 | -18.1 | **< 0.001** |

Parameter Estimates for the Fixed Effects with meaningful and anomalous as the reference contrasts for the pairwise comparisons in the Linear Mixed-Effects Models. m = meaningful, a = anomalous, p = pseudowords.

**Activation Tables**

Activation maps were thresholded at p < .05 FDR-corrected. Up to 5 peaks per cluster that are more than 8 mm apart are reported.

For the psychophysiological interaction, Contrast images were thresholded at p < 0.05, cluster-level family wise error (FWE) corrected, with a voxel-wise threshold of p < 0.001. L = left, R = right, ACC = anterior cingulate cortex, AG = angular gyrus, ATL = anterior temporal lobe, IFG = inferior frontal gyrus, ITG = inferior temporal gyrus, IPS = intraparietal sulcus, SMA = supplementary motor area, SMG = supramarginal gyrus, M1 = primary motor cortex, MTG = middle temporal gyrus, PFC = prefrontal cortex, MCC = middle cingulate cortex, v = ventral, d = dorsal, m = medial, p = posterior, a = anterior.

**Supplementary Table 3.** Activation peaks for meaningful > pseudoword phrases in the implicit task

| **Region** | **Cluster size (voxel/mm³)** | **x** | **y** | **z** | **T** |
| --- | --- | --- | --- | --- | --- |
| **L AG** | 262/4093 |  |  |  |  |
| L AG (PGp) |  | -50 | -70 | 28 | 6.91 |
| L AG |  | -42 | -57 | 18 | 5.89 |
| L AG (PGp) |  | -52 | -67 | 20 | 5.41 |
| **L DMPFC** | 62/968 |  |  |  |  |
| L middle frontal gyrus |  | -30 | 26 | 42 | 5.05 |
| L superior frontal gyrus |  | -20 | 38 | 42 | 4.65 |
| L superior frontal gyrus |  | -12 | 43 | 45 | 4.54 |
| **L pITG/pMTG** | 34/531 |  |  |  |  |
| L pITG |  | -52 | -30 | -18 | 4.94 |
| L pMTG |  | -62 | -22 | -15 | 4.74 |
| **R ACC** | 23/369 |  |  |  |  |
| R ACC |  | 3 | 30 | -5 | 4.92 |

**Supplementary Table 4.** Activation peaks for meaningful > pseudoword phrases in the explicit task

| **Region** | **Cluster size (voxels/mm ³)** | **x** | **y** | **z** | **T** |
| --- | --- | --- | --- | --- | --- |
| **L aIFG, DMPFC, vmPFC** | 6400/100000 |  |  |  |  |
| L middle orbital gyrus |  | -47 | 48 | -8 | 7.65 |
| L aIFG (pars orbitalis) |  | -47 | 40 | -12 | 7.05 |
| L aIFG (pars orbitalis) |  | -34 | 36 | -12 | 6.88 |
| L aIFG (pars orbitalis) |  | -32 | 20 | -18 | 6.79 |
| L superior medial gyrus |  | -2 | 46 | 42 | 6.61 |
| **L AG, pSMG, IPS** | 1879/29359 |  |  |  |  |
| L AG (PGp) |  | -40 | -80 | 32 | 6.59 |
| L AG (PGp) |  | -47 | -70 | 28 | 6.22 |
| L AG (PGa) |  | -44 | -60 | 42 | 5.78 |
| L AG |  | -40 | -57 | 25 | 5.58 |
| L AG (PGp) |  | -52 | -64 | 20 | 5.23 |
| **L pMTG/ITG/ATL** | 1186/18531 |  |  |  |  |
| L pMTG |  | -57 | -50 | -8 | 7.35 |
| L pITG |  | -60 | -42 | -12 | 7.27 |
| L pITG |  | -54 | -57 | -12 | 7.01 |
| L pMTG |  | -64 | -32 | -15 | 5.43 |
| L ATL |  | -52 | -2 | -38 | 5.12 |
| **R cerebellum** | 887/13859 |  |  |  |  |
| R cerebellum (crus I) |  | 40 | -74 | -42 | 6.72 |
| R cerebellum (crus II) |  | 18 | -84 | -38 | 6.02 |
| R cerebellum (crus I) |  | 13 | -80 | -25 | 5.84 |
| R cerebellum (crus I) |  | 36 | -77 | -35 | 5.5 |
| R cerebellum (crus I) |  | 43 | -72 | -35 | 5.43 |
| **R M1, premotor, somatosensory cortex** | 274/4812 |  |  |  |  |
| R primary somatosensory cortex |  | 36 | -27 | 48 | 4.28 |
| R primary motor cortex |  | 38 | -20 | 50 | 4.17 |
| R primary motor cortex |  | 33 | -12 | 52 | 3.88 |
| R primary somatosensory cortex |  | 48 | -22 | 60 | 3.59 |
| R primary somatosensory cortex |  | 50 | -17 | 48 | 3.54 |
| **R insula, temporal pole** | 219/3421 |  |  |  |  |
| R insula |  | 38 | 23 | -8 | 6.4 |
| R temporal pole |  | 40 | 10 | -20 | 4.33 |
| R temporal pole |  | 28 | 8 | -20 | 3.87 |
| R temporal pole |  | 30 | 3 | -12 | 3.26 |
| **L/R ACC** | 128/2000 |  |  |  |  |
| L ACC |  | -4 | 6 | 28 | 4.54 |
| R ACC |  | 8 | 8 | 28 | 4.3 |
| **R cerebellum** | 109/1703 |  |  |  |  |
| R cerebellum (lobule IX) |  | 3 | -57 | -48 | 5.44 |
| R cerebellum (lobule IX) |  | 10 | -50 | -40 | 4.98 |
| R cerebellum (lobule IX) |  | 20 | -47 | -40 | 3.68 |
| **L PCC/MCC** | 87/1359 |  |  |  |  |
| L PCC |  | -2 | -42 | 32 | 3.64 |
| L MCC |  | -7 | -27 | 38 | 3.5 |
| L MCC |  | 0 | -34 | 35 | 3.38 |
| **brain stem/thalamus** | 77/1203 |  |  |  |  |
| brain stem |  | -2 | -34 | -35 | 4.63 |
| brain stem |  | 0 | -24 | -35 | 3.36 |
| brain stem |  | -10 | -40 | -40 | 3.3 |
| brain stem |  | 13 | -30 | -35 | 3.26 |
| brain stem |  | 6 | -27 | -42 | 2.93 |
| **L cerebellum** | 57/890 |  |  |  |  |
| L cerebellum (crus II) |  | -34 | -74 | -40 | 4.16 |
| L cerebellum (crus I) |  | -27 | -70 | -32 | 3.39 |
| **R cerebellum** | 55/859 | 3 | -57 | -28 | 4.43 |
| **L precuneus** | 52/812 | -2 | -54 | 12 | 4.61 |
| **L insula** | 46/718 |  |  |  |  |
| L insula |  | -42 | -2 | 8 | 4.27 |
| L insula |  | -40 | -4 | 0 | 3.39 |
| **R fusiform, cerebellum** | 45/703 |  |  |  |  |
| R cerebellum (lobule IV-V) |  | 23 | -32 | -22 | 3.62 |
| R fusiform gyrus |  | 28 | -42 | -18 | 3.6 |
| R parahippocampal gyrus |  | 36 | -37 | -12 | 3.19 |
| **R posterior-medial frontal gyrus** | 43/671 | 10 | -7 | 55 | 4.55 |
| **L hippocampus** | 42/656 |  |  |  |  |
| L hippocampus |  | -30 | -12 | -12 | 4.29 |
| **R fusiform gyrus** | 41/640 |  |  |  |  |
| R fusiform gyrus |  | 38 | -17 | -25 | 4.36 |
| **R pITG** | 29/453 |  |  |  |  |
| R pITG |  | 56 | -47 | -12 | 3.44 |
| **R insula** | 26/406 |  |  |  |  |
| R insula |  | 40 | -7 | 0 | 3.52 |
| R insula |  | 40 | 6 | 0 | 3.22 |
| **R middle orbital gyrus** | 23/359 | 33 | 40 | -10 | 4 |
| **R AG (PGp)** | 23/359 | 53 | -70 | 25 | 3.62 |

**Supplementary Table 5**. Activation peaks for anomalous > pseudoword phrases in the explicit task

| **Region** | **Cluster size (voxels/mm ³)** | **x** | **y** | **z** | **T** |
| --- | --- | --- | --- | --- | --- |
| **L aIFG, DMPFC** | 3690/57656 |  |  |  |  |
| L aIFG (pars orbitalis) |  | -42 | 28 | -12 | 8.86 |
| L posterior-medial frontal gyrus |  | -2 | 18 | 50 | 8.71 |
| L aIFG (pars triangularis) |  | -52 | 36 | 10 | 7.91 |
| L aIFG (pars orbitalis) |  | -37 | 36 | -15 | 7.48 |
| L superior medial gyrus |  | -10 | 38 | 50 | 7.28 |
| **R cerebellum** | 679/10609 |  |  |  |  |
| R cerebellum (crus I) |  | 36 | -72 | -42 | 7.21 |
| R cerebellum (crus II) |  | 26 | -74 | -48 | 6.21 |
| R cerebellum (crus I) |  | 18 | -77 | -28 | 6.11 |
| R cerebellum (crus II) |  | 23 | -82 | -42 | 5.56 |
| R cerebellum (crus I) |  | 28 | -72 | -28 | 4.33 |
| **L/R thalamus, caudate nucleus** | 422/6593 |  |  |  |  |
| R caudate nucleus |  | 13 | 8 | 12 | 6.07 |
| L thalamus |  | -4 | -22 | 8 | 5.3 |
| L thalamus |  | -7 | -12 | 8 | 4.9 |
| L caudate nucleus |  | -12 | 10 | 10 | 4.86 |
| L thalamus |  | -12 | -2 | 12 | 4.78 |
| **L MTG/ITG** | 365/5703 |  |  |  |  |
| L pMTG |  | -62 | -47 | -2 | 6.17 |
| L pMTG |  | -50 | -37 | -2 | 5.34 |
| L pITG |  | -52 | -52 | -10 | 4.6 |
| L pITG |  | -50 | -62 | -10 | 4.14 |
| L pMTG |  | -60 | -62 | 2 | 3.18 |
| **L AG/SMG** | 225/3515 |  |  |  |  |
| L AG |  | -40 | -57 | 25 | 5.7 |
| L AG (PGp) |  | -44 | -70 | 25 | 5.3 |
| **L fusiform gyrus** | 164/2562 |  |  |  |  |
| L fusiform gyrus |  | -32 | -30 | -25 | 6.1 |
| L fusiform gyrus |  | -42 | -17 | -22 | 5.39 |
| L fusiform gyrus |  | -40 | -27 | -18 | 4.95 |
| **L ATL, temporal pole, parahippocampal gyrus** | 137/2140 |  |  |  |  |
| L ATL |  | -47 | -4 | -32 | 5.62 |
| L temporal pole |  | -32 | 6 | -42 | 4.64 |
| L ATL |  | -42 | -4 | -40 | 4.06 |
| L temporal pole |  | -47 | 10 | -30 | 3.82 |
| L temporal pole |  | -42 | 3 | -32 | 3.62 |
| **R aIFG, insula** | 133/2078 |  |  |  |  |
| R insula |  | 36 | 20 | -8 | 5.25 |
| R insula |  | 33 | 26 | 2 | 3.77 |
| R aIFG (pars orbitalis) |  | 28 | 26 | -12 | 3.66 |
| R aIFG (pars orbitalis) |  | 43 | 28 | -12 | 3.11 |
| **R aIFG** | 51/796 |  |  |  |  |
| R aIFG (pars orbitalis) |  | 36 | 38 | -8 | 4.84 |
| R aIFG (pars orbitalis) |  | 48 | 40 | -12 | 3.68 |
| **R amygdala** | 41/640 |  |  |  |  |
| R amygdala |  | 28 | -4 | -12 | 5.12 |
| R pallidum |  | 20 | -2 | -2 | 3.42 |
| **L thalamus** | 32/500 |  |  |  |  |
| L thalamus |  | -30 | -14 | -8 | 3.83 |
| L thalamus |  | -24 | -22 | -8 | 3.17 |
| **brainstem** | 27/421 |  |  |  |  |
| brainstem |  | -4 | -27 | -25 | 4.57 |
| **L ATL** | 25/390 |  |  |  |  |
| L ATL |  | -54 | 3 | -20 | 4.3 |
| **R cerebellum** | 22/343 |  |  |  |  |
| R cerebellum (lobule IX) |  | 3 | -57 | -50 | 4.19 |
| R cerebellum (lobule IX) |  | 8 | -64 | -42 | 3.48 |
| **R cerebellum** | 22/343 |  |  |  |  |
| R cerebellum (crus II) |  | -32 | -77 | -42 | 3.59 |
| **R thalamus** | 21/328 |  |  |  |  |
| R thalamus |  | 18 | -20 | 0 | 4.64 |

**Supplementary Table 6.** Activation peaks for conjunction of [explicit: meaningful > pseudowords] ∩ [explicit: anomalous > pseudowords]

| **Region** | **Cluster size (voxels/mm³)** | **x** | **y** | **z** | **T** |
| --- | --- | --- | --- | --- | --- |
| **L aIFG, DMPFC** | 2679/41859 |  |  |  |  |
| L aIFG (pars orbitalis) |  | -47 | 40 | -12 | 7.05 |
| L aIFG (pars orbitalis) |  | -34 | 33 | -12 | 6.58 |
| L superior medial gyrus |  | -12 | 38 | 50 | 6.39 |
| L superior medial gyrus |  | -14 | 36 | 55 | 6.28 |
| L aIFG (pars orbitalis) |  | -47 | 46 | -5 | 6.24 |
| **R cerebellum** | 523/8171 |  |  |  |  |
| R cerebellum (Crus II) |  | 38 | -74 | -42 | 6.52 |
| R cerebellum (Crus I) |  | 13 | -80 | -25 | 5.8 |
| R cerebellum (Crus II) |  | 18 | -87 | -38 | 4.74 |
| R cerebellum (Crus II) |  | 20 | -84 | -40 | 4.71 |
| R cerebellum (VIII) |  | 26 | -74 | -48 | 4.28 |
| **L/R thalamus, caudate nucleus** | 357/5578 |  |  |  |  |
| L caudate nucleus |  | -12 | 10 | 10 | 4.86 |
| L caudate nucleus |  | -12 | 0 | 12 | 4.63 |
| L thalamus |  | -2 | -22 | 8 | 4.42 |
| L caudate nucleus |  | -10 | 13 | 2 | 4.31 |
| R caudate nucleus |  | 13 | 10 | 10 | 4.18 |
| **L pMTG/ITG** | 333/5203 |  |  |  |  |
| L pMTG |  | -62 | -47 | -2 | 6.17 |
| L pITG |  | -52 | -52 | -10 | 4.6 |
| L pITG |  | -54 | -57 | -8 | 4.32 |
| L pITG |  | -50 | -62 | -10 | 4.14 |
| L pMTG |  | -60 | -62 | 2 | 3.18 |
| **L AG/SMG** | 224/3500 |  |  |  |  |
| L AG |  | -40 | -57 | 25 | 5.58 |
| L SMG (PFm) |  | -42 | -60 | 22 | 5.56 |
| L AG (PGp) |  | -44 | -70 | 25 | 5.3 |
| **L ATL** | 112/1750 |  |  |  |  |
| L ATL |  | -47 | -4 | -32 | 4.69 |
| L ATL |  | -32 | 3 | -40 | 3.72 |
| L ATL |  | -44 | 8 | -32 | 3.59 |
| L ATL |  | -30 | 0 | -35 | 3.09 |
| L pITG |  | -50 | 10 | -30 | 3.08 |
| **L ITG/fusiform** | 102/1593 |  |  |  |  |
| L fusiform gyrus |  | -30 | -32 | -20 | 5.29 |
| L fusiform gyrus |  | -32 | -30 | -22 | 5.19 |
| L fusiform gyrus |  | -27 | -40 | -18 | 4.04 |
| L fusiform gyrus |  | -42 | -24 | -20 | 3.66 |
| **R insula/IFG** | 83/1296 |  |  |  |  |
| R Insula |  | 36 | 20 | -8 | 5.25 |
| R IFG (pars orbitalis) |  | 36 | 23 | -15 | 3.41 |

**Supplementary Table 7**. Activation peaks for meaningful > anomalous phrases in the explicit task

| **Region** | **Cluster size (voxels/mm³)** | **x** | **y** | **z** | **T** |
| --- | --- | --- | --- | --- | --- |
| **L and R ACC, DMPFC** | 328/ 5125 |  |  |  |  |
| L ACC |  | -2 | 48 | 5 | 5.91 |
| L superior medial gyrus |  | 0 | 58 | 10 | 5.42 |
| L ACC |  | -2 | 36 | 8 | 4.9 |
| L superior medial gyrus |  | 0 | 46 | 20 | 4.84 |
| R ACC |  | 3 | 33 | 15 | 4.5 |
| **L AG, IPS, SMG (PF/PFm)** | 286/4468 |  |  |  |  |
| L AG (PGa/PFm) |  | -44 | -60 | 42 | 5.59 |
| L SMG (PF) |  | -57 | -42 | 48 | 5.36 |
| L AG (PGa) |  | -44 | -54 | 55 | 5.26 |
| L IPS (hIP3) |  | -34 | -50 | 50 | 4.88 |
| L AG (PGa) |  | -42 | -70 | 42 | 4.72 |
| **L pMTG** | 90/1406 |  |  |  |  |
| L pMTG |  | -60 | -32 | -15 | 5.11 |
| L pMTG |  | -60 | -52 | -8 | 4.45 |
| L pMTG |  | -57 | -44 | -10 | 4.17 |
| **L vmPFC** | 82/1281 |  |  |  |  |
| L superior orbital gyrus |  | -12 | 18 | -20 | 6.25 |
| L rectal gyrus (s32) |  | -7 | 28 | -18 | 5 |
| L rectal gyrus |  | -14 | 30 | -15 | 4.53 |
| **R AG** | 42/656 |  |  |  |  |
| R AG (PGp) |  | 40 | -70 | 38 | 5.19 |
| R middle occipital gyrus |  | 43 | -77 | 32 | 4.68 |
| **R caudate nucleus** | 24/375 |  |  |  |  |
| R caudate nucleus |  | 8 | 13 | 8 | 6.14 |
| **L MCC** | 20/312 |  |  |  |  |
| L MCC |  | -4 | -22 | 40 | 4.36 |

**Supplementary Table 8**. Activation peaks for conjunction of [explicit: meaningful > pseudowords] & [implicit: meaningful > pseudowords]

| **Region** | **Cluster size (voxel/mm³)** | **x** | **y** | **z** | **T** |
| --- | --- | --- | --- | --- | --- |
| **AG (PGp), pSMG** | 259/4046 |  |  |  |  |
| L AG (PGp) |  | -47 | -70 | 28 | 6.22 |
| L AG (PGp) |  | -40 | -74 | 28 | 5.23 |
| L AG (PGp) |  | -52 | -64 | 20 | 5.01 |
| L AG |  | -42 | -57 | 22 | 4.9 |
| L SMG (PFcm) |  | -50 | -47 | 30 | 4.35 |
| **L pITG** | 21/328 |  |  |  |  |
| L pITG |  | -60 | -37 | -20 | 3.54 |
| L pITG |  | -52 | -32 | -18 | 3.27 |
| **L DMPFC** | 20/312 |  |  |  |  |
| L superior frontal gyrus |  | -17 | 43 | 42 | 4.6 |
| L superior frontal gyrus |  | -12 | 43 | 45 | 4.53 |

**Supplementary Table 9**. Interaction: Explicit > Implicit task for meaningful > pseudoword phrases (inclusively masked with significant voxels from the explicit task: meaningful > pseudoword).

| **Region** | **Cluster size (voxels/mm³)** | **x** | **y** | **z** | **T** |
| --- | --- | --- | --- | --- | --- |
| **L aIFG, insula** | 1132/17688 |  |  |  |  |
| L insula |  | -32 | 20 | 0 | 9.12 |
| L IFG (pars triangularis) |  | -52 | 26 | 25 | 6.69 |
| L insula |  | -27 | 20 | -10 | 6.26 |
| L IFG (pars triangularis) |  | -47 | 33 | 18 | 6.13 |
| L precentral gyrus |  | -40 | 6 | 42 | 6.08 |
| **L/R thalamus, caudate nucleus** | 899/14047 |  |  |  |  |
| R thalamus |  | 13 | -7 | 2 | 5.81 |
| R caudate nucleus |  | 13 | 10 | 8 | 5.75 |
| L caudate nucleus |  | -10 | 13 | 5 | 5.71 |
| L thalamus |  | -10 | -12 | 2 | 5.4 |
| R thalamus |  | 6 | -27 | 5 | 5.27 |
| **L/R ACC, DMPFC** | 697/10891 |  |  |  |  |
| L superior medial frontal gyrus |  | -4 | 28 | 42 | 7.13 |
| R superior medial frontal gyrus |  | 3 | 30 | 38 | 7.02 |
| L superior medial frontal gyrus |  | -2 | 18 | 48 | 6.97 |
| R superior medial frontal gyrus |  | 6 | 23 | 42 | 6.47 |
| L posterior-medial frontal gyrus |  | -2 | 23 | 60 | 5.53 |
| **L IPS/SMG** | 445/6953 |  |  |  |  |
| L SMG (PFt) |  | -44 | -37 | 48 | 6.4 |
| L IPS (hIP3) |  | -40 | -47 | 45 | 6.11 |
| L SMG (PFm) |  | -52 | -60 | 42 | 4.15 |
| L IPS (hIP3) |  | -32 | -60 | 42 | 3.86 |
| L SMG (PFt) |  | -52 | -34 | 45 | 3.85 |
| **L pMTG/ITG** | 227/3547 |  |  |  |  |
| L pITG |  | -47 | -47 | -12 | 4.57 |
| L pITG |  | -57 | -52 | -12 | 4.49 |
| L pITG |  | -57 | -42 | -12 | 4.05 |
| L pMTG |  | -64 | -44 | -5 | 3.45 |
| L pMTG |  | -54 | -40 | -5 | 3.13 |
| **R cerebellum** | 196/3063 |  |  |  |  |
| R cerebellum (crus I) |  | 26 | -67 | -30 | 4.4 |
| R cerebellum (crus II) |  | 26 | -72 | -48 | 4.1 |
| R cerebellum (crus II) |  | 36 | -72 | -48 | 4.07 |
| R cerebellum (crus I) |  | 48 | -62 | -32 | 3.95 |
| R cerebellum (crus I) |  | 43 | -70 | -30 | 3.81 |
| **R aIFG** | 121/1891 |  |  |  |  |
| R IFG (pars orbitalis) |  | 33 | 26 | -5 | 7.85 |
| **R cerebellum** | 78/1219 |  |  |  |  |
| R cerebellum (lobule VI) |  | 10 | -80 | -25 | 4.93 |
| R cerebellum (crus I) |  | 13 | -82 | -32 | 3.82 |
| **R postcentral** | 34/531 |  |  |  |  |
| R S1 |  | 50 | -20 | 58 | 4.24 |
| R S1 |  | 48 | -22 | 48 | 3.38 |
| **L/R ACC** | 30/469 |  |  |  |  |
| L ACC |  | -4 | 6 | 25 | 4.25 |
| R ACC |  | 6 | 8 | 28 | 3.56 |

**Supplementary Table 10.** Interaction: Explicit > implicit task for anomalous > pseudoword phrases (inclusively masked with significant voxels from the explicit task: anomalous > pseudoword).

| **Region** | **Cluster size (voxels/mm³)** | **x** | **y** | **z** | **T** |
| --- | --- | --- | --- | --- | --- |
| **L aIFG, insula** | 1186/18531 |  |  |  |  |
| L IFG (pars triangularis) |  | -52 | 33 | 10 | 6.51 |
| L insula |  | -27 | 26 | 2 | 6.46 |
| L IFG (pars triangularis) |  | -54 | 20 | 25 | 6.41 |
| L insula |  | -37 | 20 | -2 | 6.2 |
| L IFG (pars triangularis) |  | -47 | 23 | 0 | 5.92 |
| **L DMPFC** | 639/9984 |  |  |  |  |
| L posterior-medial frontal gyrus |  | -2 | 18 | 50 | 7.26 |
| L superior medial frontal gyrus |  | -2 | 28 | 42 | 5.84 |
| L posterior-medial frontal gyrus |  | 0 | 18 | 60 | 5.76 |
| L superior medial frontal gyrus |  | 0 | 36 | 50 | 4.81 |
| L superior medial frontal gyrus |  | -7 | 36 | 40 | 4.35 |
| **R cerebellum** | 141/2203 |  |  |  |  |
| R cerebellum (lobule VI) |  | 28 | -60 | -32 | 4.63 |
| R cerebellum (crus I) |  | 46 | -60 | -32 | 4.52 |
| R cerebellum (lobule VIIb) |  | 23 | -72 | -48 | 4.48 |
| R cerebellum (crus I) |  | 36 | -70 | -40 | 4.42 |
| R cerebellum (lobule VI) |  | 28 | -60 | -32 | 4.63 |
| **R thalamus/caudate nucleus** | 112/1750 |  |  |  |  |
| R thalamus |  | 13 | -17 | 12 | 5.07 |
| R caudate nucleus |  | 16 | 6 | 12 | 5.03 |
| R thalamus |  | 8 | -4 | 5 | 4.57 |
| **R aIFG** | 102/1593 |  |  |  |  |
| R IFG (pars orbitalis) |  | 33 | 26 | -5 | 7.12 |
| **R cerebellum** | 81/1265 |  |  |  |  |
| R cerebellim (lobule VI) |  | 10 | -77 | -25 | 5.41 |
| **L thalamus** | 67/1046 |  |  |  |  |
| L thalamus |  | -14 | -20 | 12 | 4.66 |
| L thalamus |  | -4 | -22 | 8 | 4.21 |
| **L caudate nucleus** | 35/546 |  |  |  |  |
| L caudate nucleus |  | -14 | 3 | 15 | 4.17 |
| L caudate nucleus |  | -12 | 8 | 8 | 4.02 |

**Supplementary Table 11.** Activation peaks for the psychophysiological interaction during the explicit task for meaningful > pseudowords seeded from the left PGp

| **Region** | **Cluster size (voxel/mm³)** | **x** | **y** | **z** | **T** |
| --- | --- | --- | --- | --- | --- |
| **Pre-SMA** | 106/1656 |  |  |  |  |
| L pre-SMA |  | -4 | 6 | 6 | 5.2 |
| R SMA |  | 8 | 0 | 60 | 4.16 |
| **L aIFG** | 73/1140 |  |  |  |  |
| L aIFG (pars triangularis) |  | -54 | 20 | 8 | 4.93 |
| L aIFG (pars triangularis) |  | -40 | 20 | 5 | 4.18 |
| L aIFG (pars orbitalis) |  | -44 | 23 | -5 | 4.03 |
